# Oscillatory responses to generated and perceived rhythms

**DOI:** 10.1101/2020.11.19.390062

**Authors:** Markus Ostarek, Phillip M. Alday, Olivia Gawel, Johannes Wolfgruber, Birgit Knudsen, Francesco Mantegna, Falk Huettig

**Affiliations:** Max Planck Institute for Psycholinguistics; Pompeu Fabra University Barcelona; Graz University of Technology; New York University; Radboud University Nijmegen

**Author notes:** Corresponding author for this preprint, Address for correspondence, Markus Ostarek), Max Planck Institute for Psycholinguistics, Wundtlaan 1, 6525 XD Nijmegen, The Netherlands.

**Keywords:** Neural oscillations, music, rhythm perception

## Abstract

Neural oscillations have been proposed as a mechanism for structure building in language and music. In music, this idea is appealing because of the intuitive mapping between perceptual and neural rhythms. The strongest evidence has come from studies in which participants listened to isochronous sequences of identical tones and were asked to imagine hearing them in binary (march) or ternary meter (waltz). The critical finding was that in addition to increased signal at the frequency corresponding to the tone rate there was increased signal at the imagined meter frequencies. While it is striking that meter tracking was observed without any acoustic cues in the input, rhythm perception was confounded with rhythm imagery involving active generation of rhythmic structure. We conducted two electroencephalography experiments with musicians and non-musicians, teasing apart the effects of rhythm perception and rhythm generation. Evidence for meter-related neural oscillations was only observed in situations where rhythmic structure was actively generated, either via rhythm imagery or in the form of overt behavior (tapping). Thus, our data suggest that mere rhythm perception is not sufficient to elicit oscillations at the meter frequency and that they are instead driven by the active generation of rhythm. This undermines the proposal that neural oscillations constitute a basic structure building mechanism in rhythm perception and raises questions about the role of oscillations in language processing.

## 1. Introduction

Humans have a unique capacity for language and music. A defining feature of both domains is that elements are combined to form higher-order chunks, giving rise to hierarchical structure. For instance, syllables are combined to form words which are combined to form phrases and sentences, and beats are combined to form bars or measures which give rise to phrases, etc. According to influential accounts, neural oscillations are a basic mechanism for hierarchical structure building in both language and music (Ding et al., 2016; Nozaradan, 2014; Nozaradan et al., 2011). For music, this view is intuitive because neural oscillations (which by definition repeat on constant time-scales) are a plausible solution for inputs that have rhythmic structure (and thus also repeat on constant time-scales).

Most of Western music uses 4/4 or 3/4 time where a new bar begins every four or three beats, making the beginning of each bar highly predictable. There is evidence (Nozaradan, 2014; Nozaradan et al., 2011, 2012, 2013; Tal et al., 2017) suggesting that sequences of isochronous tones/blips lead to enhanced power at the frequency of the tones (e.g., if there are 4 tones per second, a 4 Hz peak is observed in the power spectrum). It is still debated whether such effects can be accounted for by evoked potentials reflecting repeated exogenous acoustic stimulation (Capilla et al., 2011; Doelling et al., 2019). In line with the view that endogenous oscillations (rhythmic fluctuations in excitability which do not simply reflect evoked responses to acoustic events) underlie these effects, it has been demonstrated that beta power oscillates along with isochronous tones in an anticipatory fashion and remains high when expected tones are withheld (Fujioka et al., 2009, 2012, 2012). Moreover, a study recently reported increased phase-locking to missing beats when a feeling of pulse is generated by syncopated rhythms (Tal et al., 2017), which cannot be a direct result of exogenous stimulation because no acoustic stimulus was present at the missing beats. Finally, there is evidence that perception is cyclically modulated in silent gaps after rhythmic stimulation, suggesting that oscillations entrain to the stimulus frequency and persist after stimulus onset (De Graaf et al., 2013; Fiebelkorn et al., 2011; Spaak et al., 2014; VanRullen & Macdonald, 2012).

According to neural resonance theory (Large et al., 2015), neural oscillations can also account for the perception of musical meter. Following Patel (2006), we define meter as the “hierarchical organization of beats in which some beats are perceived as stronger than others” (p. 100). In this context, we use the term “rhythm perception” to refer to the ability to perceive acoustic events as having metric structure. Recent studies provided evidence for a link between perception of musical meter and neural entrainment by showing that sequences of identical tones (e.g., displayed at a rate of 2.4 Hz) can lead to differential neural responses depending solely on the metric interpretation of the listener (Fujioka et al., 2010, 2012, 2015; Li et al., 2019; Nozaradan et al., 2011). Using frequency tagging, recent studies observed low-frequency power peaks corresponding to the metric structure that listeners endogenously impose on the sequences (e.g., at 1.2 Hz for a binary meter and at 0.8 Hz for a ternary meter; (Li et al., 2019; Nozaradan et al., 2011)). Based on these findings, it has been proposed that simultaneous neural entrainment to beat and meter frequencies underlies rhythm perception (Nozaradan, 2014; Nozaradan et al., 2011).

Interestingly, parallel efforts in the domain of language processing have been suggested to point to the possibility that a similar mechanism could account for linguistic structure building. In a controlled setting, where linguistic (Chinese) input was engineered to be perfectly rhythmical at the level of syllables/words (4 Hz), phrases (2 Hz), and sentences (1 Hz), Ding and colleagues observed evidence for parallel neural tracking at all of these levels (Ding et al., 2016). This was only observed when the stimuli were processed by participants who understood Chinese, suggesting that acoustic bottom-up processing of the input was not responsible for the effects. Rather, the authors proposed that cortical tracking is the mechanism by which the brain builds hierarchical linguistic structure.

Thus, growing consensus appears to emerge that oscillations are a promising solution for structure building across cognitive domains. If true, this would constitute a major step for the cognitive neurosciences, as the mechanisms by which input elements are organized into larger structures have been elusive. However, there is reason for skepticism. For one thing, natural speech is not very rhythmical beyond the syllable level and it is not clear how oscillations at the frequency of phrases and sentences can account for hierarchical structure building when phrase and sentence frequencies are not stable over time (see Rimmele et al., 2018 for a related discussion of the viability of neural entrainment as a mechanism for the prediction of aperiodic stimuli). In the domain of music on which we focus here, periodicity does commonly occur but there is only limited evidence for meter tracking in the absence of meter-related acoustic cues in the input (Henry et al., 2017; Nozaradan et al., 2012, 2016). In previous studies without acoustic confounds rhythm perception was confounded with rhythm imagery (Fujioka et al., 2015; Li et al., 2019; Nozaradan et al., 2011); participants listened to isochronous tone sequences and were asked to actively imagine hearing the sequence in a binary or ternary rhythm (march or waltz). This opens up the possibility that the observed low-frequency effects at the imagined meter frequencies reflected task-specific active generation of rhythmic structure rather than rhythm perception per se.

To test whether neural entrainment to the meter frequency reflects rhythm perception, we designed two experiments. In Experiment 1, participants were cued to perceive identical tone sequences to be in 4/4 (Experiment 1a) or 3/4 time (Experiment 1b) whilst performing a secondary task unrelated to the metric structure of the sequences. This allowed us to probe the neural correlates of rhythm perception in the absence of a task that required active generation of rhythmic structure. In Experiment 2, participants were presented with the same isochronous tone sequences and were first asked to imagine hearing the tone sequence in 4/4 or 3/4 time (Experiment 2a), and then tapped along with the downbeat according to 4/4 or 3/4 time (Experiment 2b). We tested musicians and non-musicians to investigate the role of experience for oscillatory responses at the meter frequency. A recent study (Doelling & Poeppel, 2015) reported that musicians showed stronger tracking of periodically played tones (measured via inter-trial coherence) compared to non-musicians. Relatedly, a further study had participants listen to simple melodies played at a rate of 1.667 Hz and observed a relationship between musical training and cerebral acoustic coherence (Harding et al., 2019). Here, we tested whether musical training affects neural oscillations in response to perceived metric structure.

## 2. Experiment 1

Experiment 1a translated the design used by Ding et al. (2016) to the music domain, even though there cannot be complete equivalence due to differences between domains (see Figure 1). In Ding et al., mono-syllabic words were displayed at a rate of 4 Hz, with two words forming phrases and four words forming sentences. Participants were asked to monitor the sentences for swapped verb phrases that made the sentences nonsensical. In Experiment 1a, tones were presented at a rate of 4 Hz (4 tones per second) and the participants’ task was to spot occasional rhythmic irregularities. Crucially, before each tone sequence started, participants were cued with four higher pitched tones that they were told to think of as “count-in beats” that indicate the tempo of the sequence (see Palmer & Krumhansl, 1990). Ratings of perceived rhythmic structure (see Figure 2) indicated that the count-in beats had the desired effect of inducing a feeling of 4/4 time, thus giving structure to the tone sequence.

**Figure 1.**
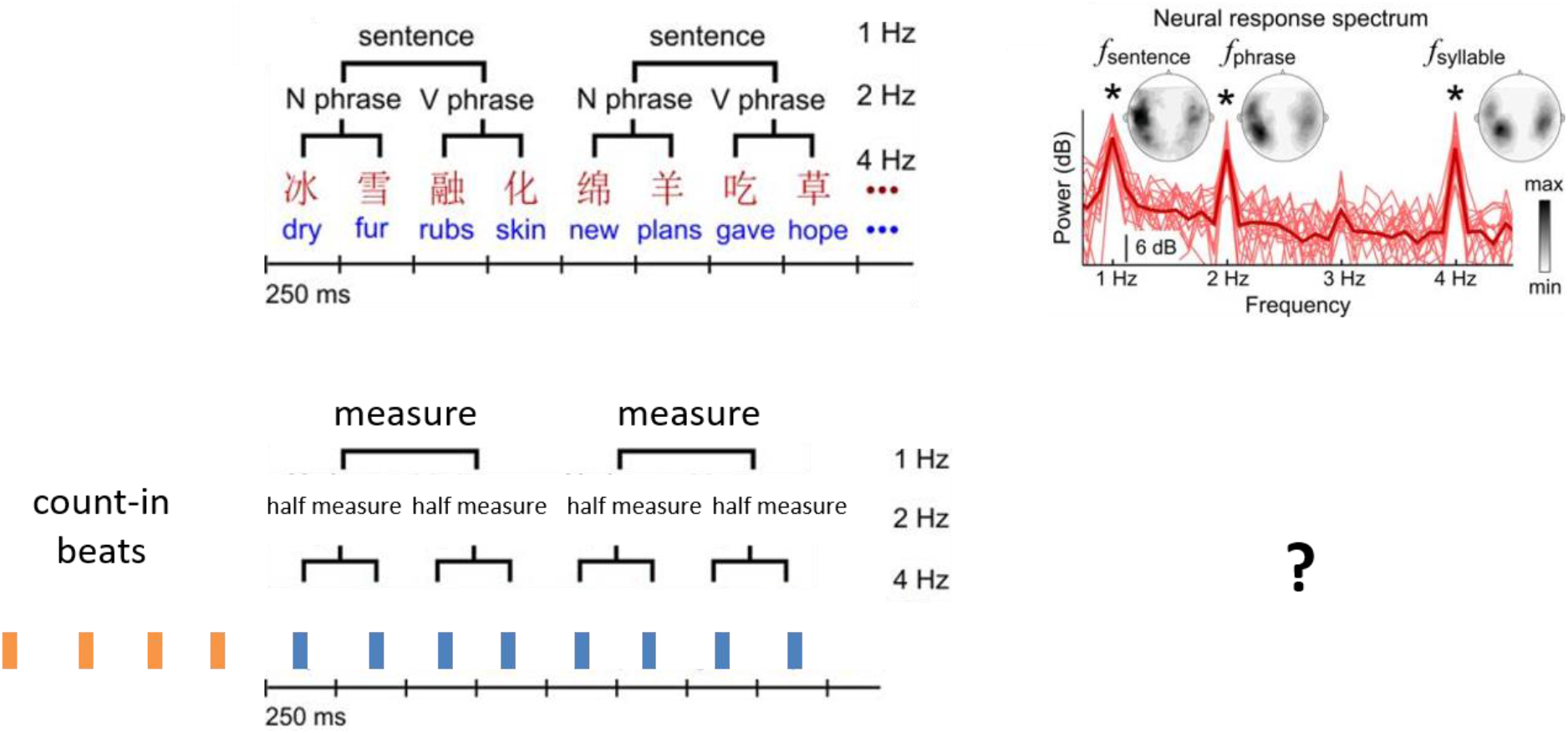
Illustration of the paradigm and its similarity to the one used by Ding et al. (2016). The main research question was whether perceiving structure in music is reflected in the pattern observed in the language domain (top right).

**Figure 2.**
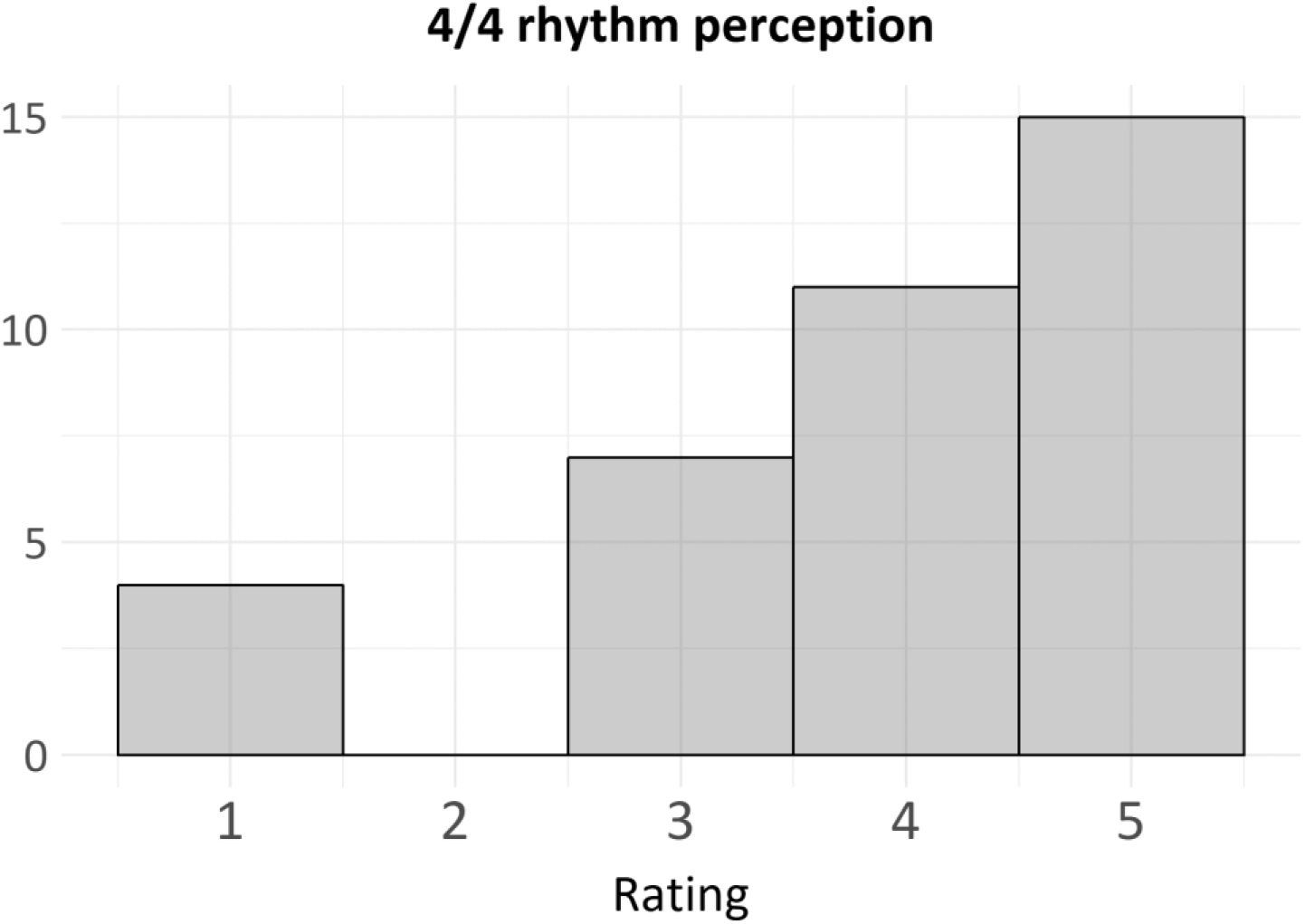
Ratings of rhythm perception collapsed across musicians and non-musicians. The y-axis shows the count per rating (1-5).

On the one hand, a clear difference between structural hierarchies in language and music is that lexical/syntactic cues were available to participants in the Ding et al. study to determine sentence structure, whereas no comparable cues were present in our minimal rhythm perception paradigm. This difference is captured by what Patel (2003) calls “syntactic representations”, which are assumed to largely differ between domains. On the other hand, it is clear that in both language and music “structural integration is a key part of syntactic processing; that is, mentally connecting each incoming element X to another element Y in the evolving structure” (Patel, 2003, p. 678). This level of “syntactic processing”, which is crucial for giving hierarchical structure to serial input elements, is a likely point of convergence between domains. With regard to this level our paradigm is similar to the one used by Ding et al. (2016) in the language domain: In both cases a rhythm (periodically occurring sentences and measures) can be perceived even though no rhythmic structure can be found in the stimulus itself. Ding et al. (2016) achieved this by stripping their stimuli of acoustic and prosodic cues which are normally available to listeners and which help constrain the interpretation of phrasal structure and syntactic dependencies (Frazier, Carlson, & Clifton, 2006). In the present study, this was similarly achieved by removing acoustic cues to structure that are typically present in music, such as regularly occurring prominent acoustic events (Hannon, Snyder, Eerola, & Krumhansl, 2004). To remove any such cues, which in our experimental setting would translate to potential confounds, we used isochronous sequences of identical tones and cued rhythmic structure before stimulus onset (see Palmer & Krumhansl, 1990), similar to the way drummers typically count in before songs begin.

Our basic design (Experiment 1a) involves stimulation at a rate of 4 Hz and cues the perception of 4/4 time, such that a new measure (rhythmic unit) is perceived to begin every four tones (i.e., at a rate of 1 Hz). The central research question was whether the perception of rhythmic structure would manifest itself in increased signal at the corresponding frequency (here 1 Hz). According to music theory (Palmer & Krumhansl, 1990), people tend to break down bars composed of four beats into two sets of two beats. Thus, an additional peak at 2 Hz might be observed. However, in the analyses below we focus on the meter frequency, as this constitutes the most direct test of our research question.

At the beginning of a subsequent EEG session, in which the data for Experiment 2 were collected, participants (n=37) were asked to rate on a scale from 1-5 to what extent they perceived 4/4 rhythm when performing this task (1=”not at all”, 5=”very strongly”). The results confirmed that both musicians and non-musicians perceived the rhythm (musicians: 3.89 (1.29), non-musicians: 3.89 (1.28)), in line with previous research showing effective cuing of perceived metric structure with count-in beats (Palmer & Krumhansl, 1990). This made it possible to test whether rhythm perception in the absence of acoustic cues and rhythm imagery leads to enhanced power at the meter frequency (1 Hz).

The musicians but not the non-musicians were invited back for Experiment 1b. Experiment 1b was identical to Experiment 1a, except that participants were cued with three count-in beats, so as to induce 3/4 time instead of 4/4 time. In this experiment, meter tracking was expected to be reflected in a peak at & ~ 1.33 Hz, as a new measure now started every 1.33s.

### 2.1 Method

#### 2.1.1 Participants

21 active musicians (10 female, mean age: 29.9, various instruments) with at least ten years of musical training and 20 non-musicians (11 female, mean age: 30.7) participated in Experiment 1a. 17 musicians (7 female, mean age: 31.6) participated in Experiment 1b. After exclusions for technical issues (three participants) and suspected intoxication (one participant), 18 musicians and 19 non-musicians entered the analysis for Experiment 1a and all 17 musicians entered the analysis for Experiment 1b. The study was covered by the Ethics Board of Radboud University. All participants gave informed consent and were paid 18 euros.

#### 2.1.2 Stimuli and Procedure

The stimuli were 30s sequences of identical tones played at a rate of four tones per second. The tones were synthetic sinusoidal tom sounds with a base frequency of 156Hz lasting 250ms with a 10ms attack time and a 240ms decay time. They were made in Ableton Live (Version 10) with the primary goal of being pleasant to listen to. Participants listened to 32 sequences, 16 of which had no irregularities, 8 had one irregularity, and 8 had two. The irregularities were produced by shifting one of the tones in the sequence forward in time by 30 ms, similar to a previous study (Nozaradan et al., 2012). The spots in the sequences where irregularities occurred varied from quite early in the sequence to very late in the sequence, to motivate participants to stay focused throughout. Before each sequence, in order to induce the perception of 4/4 or 3/4 time, four (Experiment 1a) or three (Experiment 1b) tones with higher pitch were played that participants were instructed to think of as count-in beats in music. After each trial, participants were asked to indicate whether there were zero, one, or two irregularities in the sequence, by pressing the corresponding buttons on the keyboard. Next, they were asked to initiate the following trial by pressing the space bar, thus the inter-stimulus interval was determined by the participants.

The EEG was recorded in an electrically and acoustically shielded experiment room at a sample rate of 500 Hz using an active electrode system with a BrainAmp DC amplifier (Brain Products GmbH, Gilching, Germany). We used Presentation^®^ software (Neurobehavioral Systems) for stimulus delivery and EasyCap manufactured 64-electrode montage consisting of 59 EEG channels, 4 electrooculogram (EOG), and 2 mastoid electrodes, placed according to the 10-10 system. The EOG was recorded horizontally from the electrodes placed on the left and right outer canthi and vertically from the electrodes positioned above and below the left eye. Each electrode was referenced online to the left mastoid. Electrode impedance was kept below 25 kΩ. Due to an error in recording, only 32 electrodes were recorded in some datasets. As such, we restrict the analysis in the following to the 32 electrodes available for all participants.

#### 2.1.3 EEG analyses

EEG data were processed with MNE-Python 0.17.1 (Gramfort et al., 2013). Scalp data were re-referenced to linked mastoids offline, while the EOG was re-referenced to bipolar horizontal and vertical channels. In order to eliminate ocular contributions, the EOG was then regressed out of the scalp EEG (Gratton, Coles, & Donchin 1983; Hillyard and Galambos 1970) and the signal was lowpass filtered at 10 Hz (bandpass edge, Hamming-windowed FIR, with zero-phase achieved via compensation for the group delay). The continuous EEG was then divided into epochs extending from stimulus onset (excluding the lead-in tones) until 30s post onset, thus containing the entire critical 30s interval. Epochs where the voltage exceeded 150μV were excluded from further analysis. Baseline correction was performed via mean-centering within each epoch. Only epochs without rhythmic irregularities were analyzed (n=16).

Subsequent to epoching, the complete power-spectral density (PSD) of each epoch was computed using the Fast Fourier Transform. The PSD of each epoch was then normalized by dividing by the mean power of the surrounding (±0.2 Hz) frequencies in order to remove pink noise (i.e., the tendency of spectral power to be proportional to 1/f) common in EEG (Ding et al., 2017). After normalization, the PSD was converted to the decibel scale. Due to the very precise hypotheses, we then discarded the portions of the PSD less than 0.3 Hz and more than 6 Hz.

For presentation purposes, the PSD is presented as a grand-average for each channel, i.e., averaged first across trials within each subject and then across subjects. For the statistical analysis, we only examined the frequencies reflecting our manipulation, i.e. 1 Hz (metric unit in 4/4 time), 1.33 Hz (metric unit in 3/4 time), and 4 Hz (tone rate). Experiment 1a and 1b were analyzed in the same model to assess whether the 4/4 and 3/4 time conditions differentially modulated the PSD at 1 Hz and 1.33 Hz.

Statistical analysis was performed with linear mixed-effects modeling, using the lme4 package (version 1.1.20, Bates et al. 2014) with the spectral power in decibels across all electrodes as the dependent variable. Fixed effects consisted of a categorical predictor for frequency (1 Hz vs. 1.33 Hz vs. 4 Hz), a categorical predictor for time signature (4/4 vs. 3/4), and a categorical predictor for group (musician vs. non-musician), all of which were sum-to-zero coded. Random effects consisted only of intercepts by participant. We summarize the models with Type-II Wald chi-square tests (comparable in their interpretation to traditional ANOVA) and pairwise comparisons computed via estimated-marginal means (Fox & Weisberg, 2018; Lenth, 2019).

### 2.2 Results

#### 2.2.1 Behavioral Results

Musicians (M = 90.8%, SD = 30%) were more accurate at detecting rhythmic irregularities than non-musicians (M = 78.3%, SD = 41.3%).

#### 2.2.2 EEG Results

Both in Experiment 1a and 1b, there was a strong 4 Hz peak reflecting the tone rate, but there was no evidence for the expected peaks at the meter frequencies (1 Hz peak for Experiment 1a, and 1.33 Hz for Experiment 1b), neither in musicians nor in non-musicians. There were no differences between groups (t<1). See Figure 3 for the PSD and Table 1 for the model summary of the corresponding statistical analysis.

**Figure 3.**
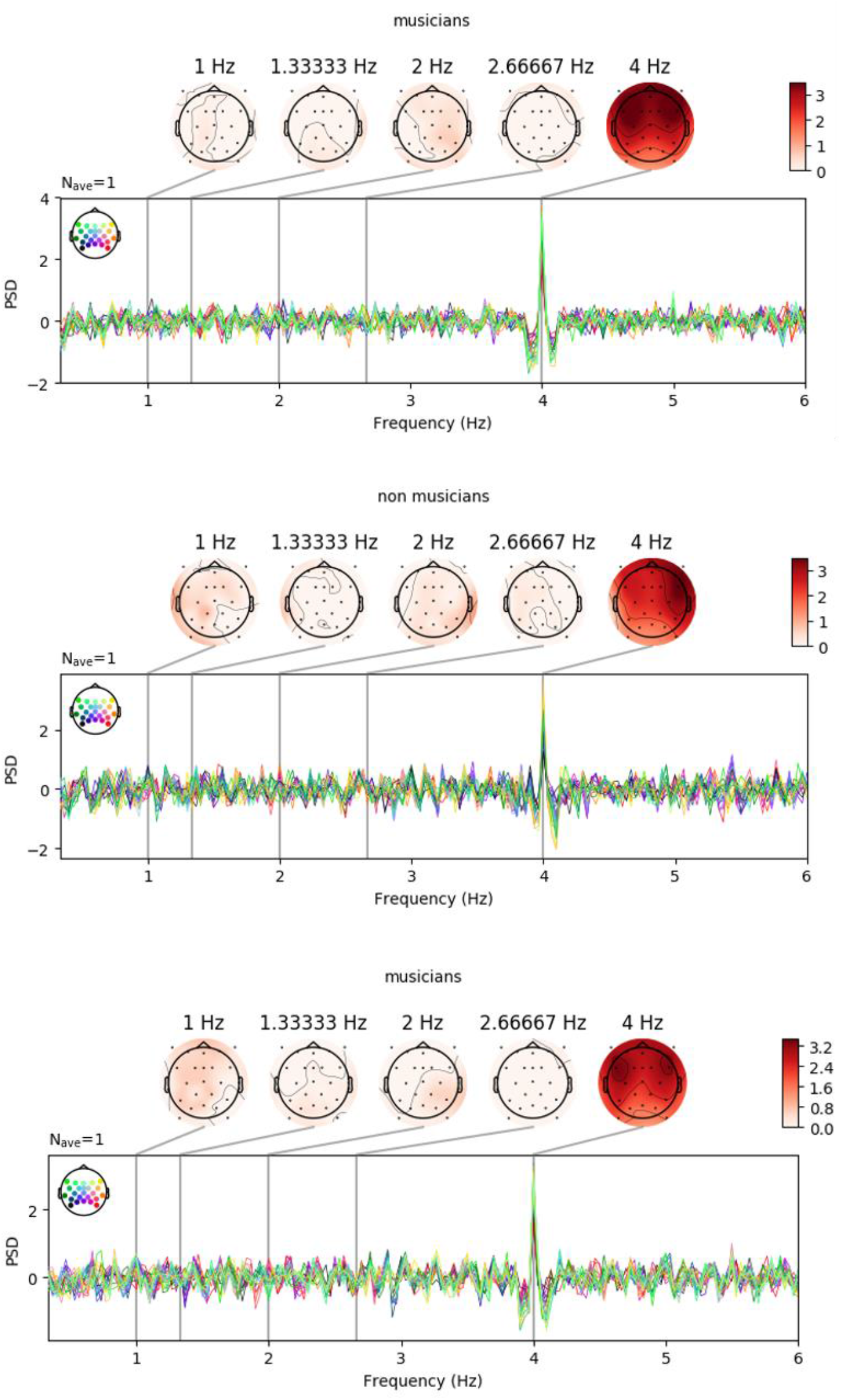
Per-channel (color-coded) power for frequencies from 1-6 Hz. The meter-related frequencies of interest (1 Hz and 1.33 Hz) and their first harmonics (2 Hz and 2.66 Hz), as well as the frequency corresponding to the stimulation rate (4 Hz) are marked with vertical lines. Top panel: Musicians in Experiment 1a. Middle panel: Non-musicians in Experiment 1a. Bottom panel: Experiment 1b (only musicians participated). Large PSD was observed at the stimulation rate (4 Hz), but not at the meter frequencies (1 Hz in Experiment 1a, 1.33 Hz in Experiment 1b) or their first harmonics (2 Hz in Experiment 1a, 2.66 Hz in Experiment 1b).

**Table 1.** Type II Wald Chi-square tests for experiment 1, collapsing across participant groups (musicians vs. non musicians). Note that there is no effect of time signature, neither as a main effect nor as an interaction.

|  | Chisq | Df | Pr(>Chisq) |
| --- | --- | --- | --- |
| frequency | 304.204 | 2 | < 0.001 |
| Time signature | 0.476 | 1 | 0.49 |
| Frequency by time signature | 1.071 | 2 | 0.582 |

**Figure 4.**
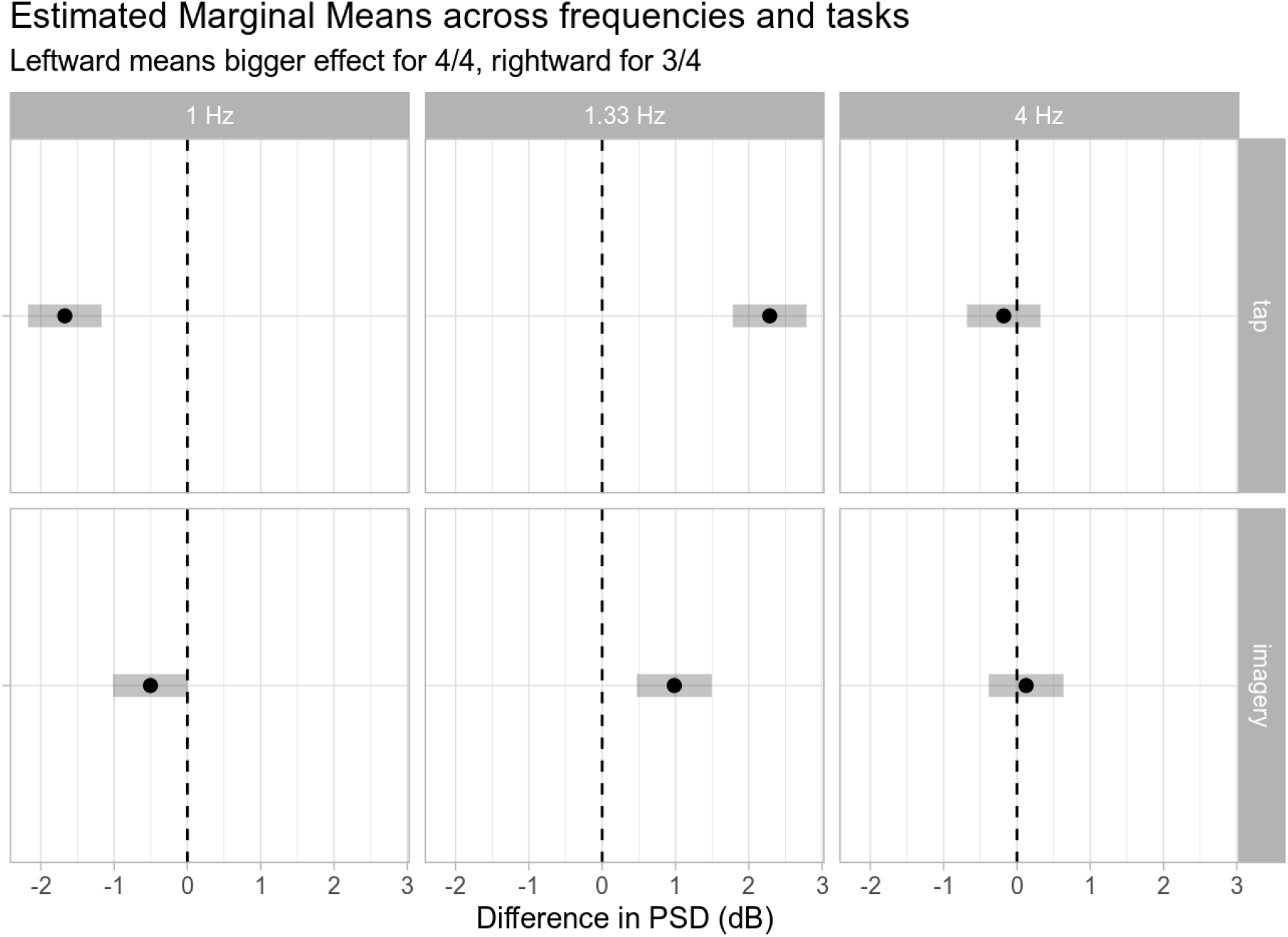
Estimated marginal means for the difference between the PSDs in the 4/4 and 3/4 time signature at different frequencies with 95% confidence intervals. Differences less than zero indicate a stronger effect in the 4/4 condition, while differences greater than zero indicate a stronger effect in the 3/4 condition. Note that the 4 Hz signal is not influenced by time signature, whereas the 1 Hz effect is enhanced in the 4/4 condition and the 1.3 Hz effect is enhanced in the 3/4 condition.

These results do not support the hypothesis that oscillations at the meter frequency play a functional role for rhythm perception, as our participants perceived the rhythm but did not show evidence for oscillations at the meter frequency in the PSD. In combination with previous findings that imagining rhythmic structure leads to robust peaks at the imagined meter frequency (Li et al., 2019; Nozaradan et al., 2011), our data suggest that such effects are related to active generation of rhythmic structure, rather than to the perception of rhythm per se. The prediction thus follows that oscillation effects, as measured by frequency tagging, should become increasingly prominent with increasingly active generation of rhythmic structure.

## 3. Experiment 2

To test the hypothesis that peaks in power at the meter frequency are driven by active generation of rhythmic structure, we modified the current paradigm slightly. Specifically, in Experiment 2a the same tone sequences (without irregularities) were presented, but now participants were asked to actively imagine hearing them in either 4/4 time (1, 2, 3, 4, 1, 2, 3, 4, …) or 3/4 time (1, 2, 3, 1, 2, 3, …). This time signature manipulation was randomized across trials. There were no more rhythmic irregularities. Before each sequence began, participants were informed which rhythm to imagine (4/4 or 3/4), and they were additionally cued with the same count-in beats as before (four vs. three higher-pitched tones). Participants were asked to rate after each trial how well they managed to imagine hearing the sequence in the cued time signature. Experiment 2a can be considered a replication of Nozaradan et al. (2011) with only slight differences regarding the sound of the tones and the number of trials. In Experiment 2b, participants listened to the same stimuli and were asked to push a button on the down beat (which would be counted as “one”), similar to (Nozaradan et al., 2013). To avoid acoustic confounds that may be introduced by sound emitted from pushing the button, we used a house-built soft button that did not make any audible noise when pushed. Before starting the tapping experiment, participants received practice until they felt comfortable with the task. In line with previous studies (Li et al., 2019; Nozaradan et al., 2011, 2013), our predictions were that we would see strong meter-related effects in the “tapping” condition and smaller but robust effects in the “imagery” condition.

### 3.1 Method

#### 3.1.1 Participants

We invited the same participants tested in Experiment 1 back to the lab. 19 musicians (8 female, mean age: 30.6) and 12 non-musicians agreed to return. To match the group sizes, eight additional non-musicians were tested to reach a total of 20 (14 female, mean age: 28.4). All participants gave informed consent and were paid 18 euros.

#### 3.1.2 EEG data collection and analysis

EEG data collection and analysis were performed the same as in Experiment 1. Additional fixed-effects were included for the task manipulation (imagery vs. tap) and for group (musician vs. non-musician). As no differences were observed between musicians and non-musicians, we collapsed across groups for the plots presented below.

### 3.2 Results

#### 3.2.1 Behavioral Results

Ratings of how well the participants managed to imagine hearing the sequence as having the cued rhythm were higher for musicians (M = 4.43, SD = 0.97) than for non-musicians (M = 3.65, SD = 1.18).

#### 3.2.2 EEG Results

As in Experiment 1, we observed strong effects at 4 Hz (the tone rate), which were not modulated by task or time signature. Crucially, both in the imagery and tap conditions, there were peaks at the meter frequencies, such that a 1 Hz peak was observed in the 4/4 condition, and a 1.33 Hz peak was observed in the 3/4 condition. As predicted, the interaction between frequency (1 Hz vs. 1.33 Hz) and time signature (4/4 vs. 3/4), was much stronger in the tap condition, but it was also present in the imagery condition (see Table 2). Note that the meter-related effects in the tapping condition do not constitute strong evidence for meter-related endogenous oscillations, but are equally compatible with periodic motor and somatosensory evoked potentials (as participants were tapping their hand periodically). There were no differences between musicians and non-musicians.

**Table 2.** Type II Wald Chi-square tests for Experiment 2. Time signature is abbreviated as & ‘ts’. Note that there are no significant effects for group, but effects for frequency, task as well as the pairwise and three-way interactions between group, frequency, and time signature.

|  | Chisq | Df | Pr(>Chisq) |
| --- | --- | --- | --- |
| frequency | 581.7762553 | 2 | 0.0000000 |
| ts | 2.8513852 | 1 | 0.0912952 |
| group | 0.0175813 | 1 | 0.8945140 |
| task | 27.5594038 | 1 | 0.0000002 |
| frequency:ts | 125.3662881 | 2 | 0.0000000 |
| frequency:group | 5.9678275 | 2 | 0.0505944 |
| ts:group | 0.0000310 | 1 | 0.9955564 |
| frequency:task | 10.6296733 | 2 | 0.0049181 |
| ts:task | 0.0609075 | 1 | 0.8050674 |
| group:task | 0.4023189 | 1 | 0.5258941 |
| frequency:ts:group | 1.3523555 | 2 | 0.5085571 |
| frequency:ts:task | 25.1396931 | 2 | 0.0000035 |
| frequency:group:task | 5.5201028 | 2 | 0.0632885 |
| ts:group:task | 0.1449769 | 1 | 0.7033826 |
| freq:ts:group:task | 0.0649615 | 2 | 0.9680411 |

Figure 5 summarizes the results of all experiments (1a, b; 2a, b) and makes the task dependence of meter-related oscillations apparent. The perception of metric structure alone does not lead to corresponding peaks in the PSD, but the stronger the task relates to the generation of metric structure, the stronger it is reflected in the PSD.

**Figure 5.**
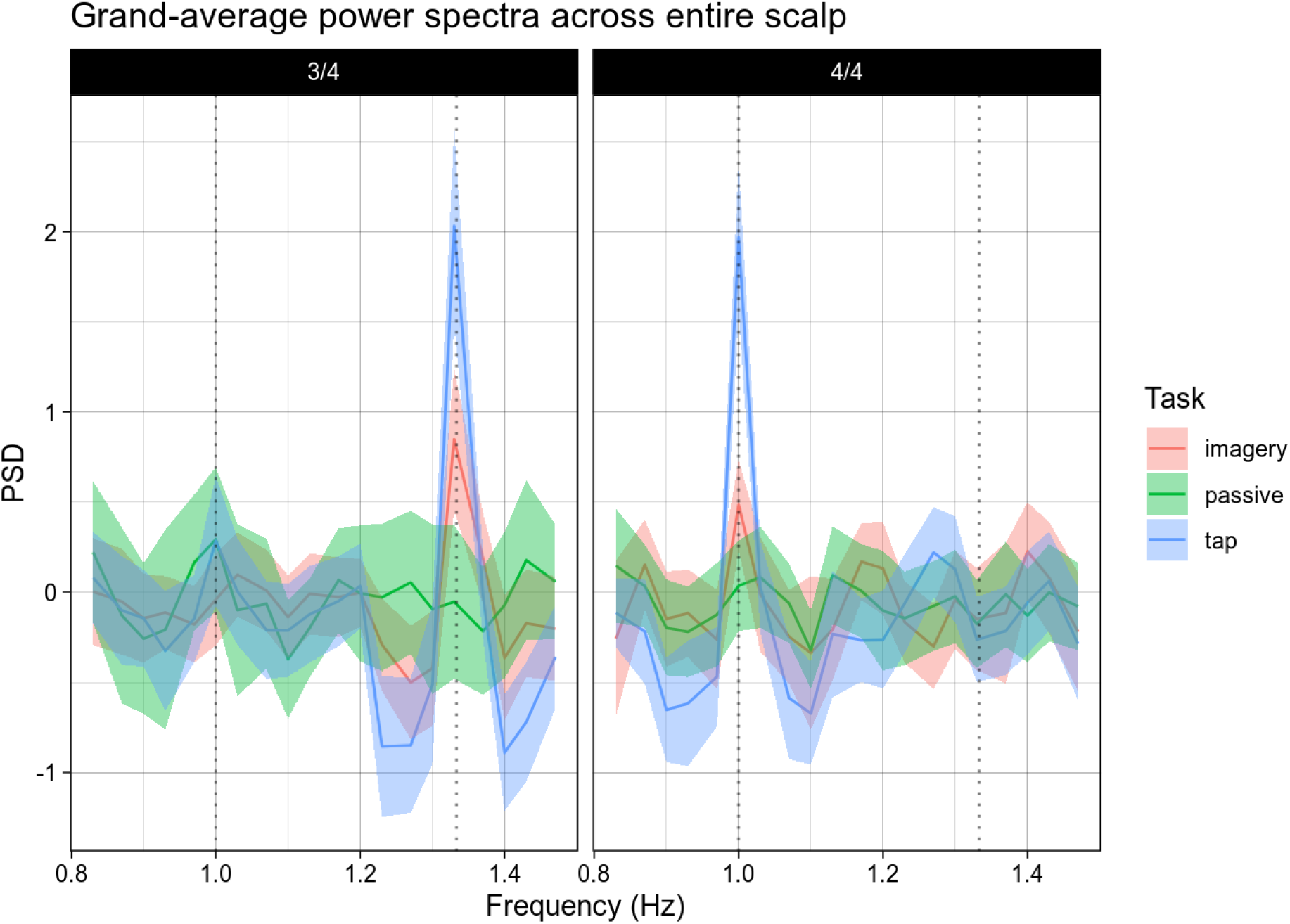
Grand-average power spectra across the entire scalp. Shaded intervals indicate 95% confidence intervals. Note that the peaks at 1.3 Hz for 3/4 and at 1 Hz for 4/4 time show a gradient structure, with the largest peaks for tapping, the second largest peaks for imagery, and no peaks for passive rhythm perception.

## 4. Discussion

Neural oscillations have been proposed as a mechanism for structure building in language and music (Ding et al., 2016; Nozaradan, 2014). In music, this idea is intuitive because of the straightforward mapping between rhythm as a perceptual/cognitive phenomenon on the one hand, and rhythm in the form of neural oscillations on the other. Neural resonance theory predicts that rhythm perception in the absence of bottom-up acoustic cues relies on low-frequency oscillations that track metric structure (Large et al., 2015). Previous evidence consistent with that theory came from studies in which rhythm perception was confounded with rhythm imagery which involves the active generation of rhythmic structure (Fujioka et al., 2010, 2012, 2015; Li et al., 2019; Nozaradan et al., 2011). Pulling apart basic rhythm perception and imagery, we observed that mere rhythm perception is not sufficient for robust meter-related frequency peaks as measured in time-domain EEG signals. Our data suggest that entrainment is driven by the active generation of rhythmic structure, either in the form of rhythm imagery or in the form of tapping.

One question that the current study left open is whether groovy rhythms rely on entrainment to the meter frequency. It is possible that in our study entrainment to tone sequences was hindered due to the stimuli’s lack of groove. In order to rule out acoustic confounds, the present paradigm used the same isochronous tone sequence across conditions. Whilst permitting maximal experimental control, this choice may come at the cost of ecological validity and rob the musical stimuli of their ability to entice the listeners. Previous studies using groovy stimuli have obtained evidence for meter-related effects (Nozaradan et al., 2012, 2016). However, it is unclear to what extent the results are driven by acoustic confounds and/or by invalid comparisons of the frequency-domain representations of the stimuli and of the neural responses (Henry et al., 2017). A recent study used a cochlear model to demonstrate that initial auditory processing differences between conditions cannot account for meter-related effects in frequency tagging analyses (Lenc et al., 2018). However, Novembre and Ianetti (2018) suggested that increased attention to salient events in syncopated patterns can give rise to effects at the meter frequency. Thus, it is unclear whether frequency tagging effects in response to groovy stimuli reflect rhythm perception, attention to periodically occurring salient events, or a mix of both. This debate indicates that it is difficult (even though it may be possible) to rid syncopated groovy stimuli of confounds that complicate the interpretation of meter-related frequency-domain effects. As of now, using isochronous sequences appears to be the safest choice as any differences are likely due to endogenously imposed rhythmic structure. It will be a challenge for future studies to strike a good balance between ecological validity and experimental control to further delineate the situations in which metric structure is tracked by neural oscillations.

Our data suggest that oscillations at the meter frequency are not responsible for the ability to perceive metric structure. This raises the question of what neural mechanism is. Previous research has demonstrated that the perception of an acoustic event is modulated by whether it is perceived to be strong or weak. It has long been known that subjective accents can be imposed on sequences of identical tones played at a fixed rate. They can be induced via rhythm imagery and have been reported to come with increased neural responses (Fujioka et al., 2010; Schaefer et al., 2011). The perception of alternating strong and weak beats even occurs even without a metric cue or imagery. This phenomenon is often referred to as the “tick-tock” phenomenon, as most people have experienced it when listening to the ticks (and tocks) of a clock. In the context of spontaneous (non-induced) subjective accenting, there is evidence for differential ERPs to tones at strong vs. weak positions in a paradigm where deviant tones were inserted in isochronous sequences (Brochard et al., 2003; Potter et al., 2009). Thus, it is possible that metric structure is maintained by up-regulating neural responses to metrically strong vs. weak beats, which could naturally account for the perceptual differences that are commonly observed for different metric positions.

The present result that active generation of rhythmic structure, but not rhythm perception, results in meter-related frequency peaks, poses a puzzle: In the language domain, sentence-level neural tracking has been observed in a context where participants were not explicitly asked to generate a “sentence rhythm” (Ding et al., 2016). Thus, one intriguing possibility is that language and music differ in the way that entrainment is deployed. It is conceivable that language is such a fundamental and important cognitive function that relevant units, such as sentences, are more naturally tracked than structural units in music. By that interpretation, one simply cannot help tracking hierarchical linguistic structure, from syllables to words, to phrases and sentences. This is consistent with the strong resilience of speech perception to noise in the environment (Ding & Simon, 2013).

However, there are strong arguments against this interpretation: 1) as noted in the Introduction, language is not usually rhythmic beyond the syllable level. With sentence length varying considerably in speech (Wiggers & Rothkrantz, 2007), it is implausible that oscillations at the “sentence frequency” play a crucial role in language comprehension, since a stable sentence rhythm is not commonly present in speech (see Rimmele et al., 2018 for arguments on why neural entrainment is not a viable mechanism for tracking non-regular events). It is likely that artificially inducing a sentence rhythm is highly noticeable to participants and makes them anticipate and “check off” the onsets of new sentences and phrases, resulting in the observed frequency peaks in the Ding et al. study. This is not expected to happen in typical language processing. Note that even if people commonly predict sentence on/offsets in natural speech this would not result in power increases at a specific frequency (band) because sentences vary too much in length. 2) In a recent study on the role of attention for neural tracking of linguistic structure, passive watching of a silent movie made robust word tracking vanish (Ding et al., 2018). As watching a silent movie without an additional task is not very demanding, one can assume that participants were still processing the words embedded in streams of syllables, but no oscillation effect was observed because participants were not actively tracking and predicting the occurrence of words across time.

Hence, oscillations may have similar functions across domains: active tracking of rhythmically repeating structure. By that interpretation, oscillatory tracking can be viewed as a task-dependent index of cognitive processes that can optionally be activated in addition to those necessary for rhythm perception and language comprehension. As these two interpretations (oscillations as a central mechanism vs. oscillations as an index of optional processes not necessary for music perception and language comprehension per se) differ quite strongly, it would be critical for future research to further test them.

Interestingly, we did not observe differences between musicians and non-musicians at the neural level (as measured with EEG). While this is not the first study in the domain of rhythm perception that obtained such a pattern (Geiser et al., 2009), other studies observed robust differences: the inter-trial coherence in response to isochronous tone sequences at a rate of ca. 1 Hz was higher in musicians (Doelling & Poeppel, 2016), an earlier P3 was observed in musicians in a deviant detection task (Ungan et al., 2013), only musicians showed a mismatch negativity when strings of tones could be grouped by good continuation of pitch (Zuijen et al., 2004), and another study found a reduced mismatch negativity in musicians performing a deviant detection task (Zhao et al., 2017). On the one hand, these results point to potential markers of music experience. On the other hand, the various measures that have been used make it difficult to evaluate how robust any single candidate marker is. High-powered pre-registered studies will be crucial to shed light on this issue.

In sum, our results challenge the view that neural oscillations at the meter frequency constitute or reflect a mechanism for rhythm perception. Instead, they suggest that meter-related oscillations reflect overt or covert generation of rhythmic structure.

## Acknowledgements

We would like to thank Nai Ding and David Poeppel for their feedback in the conception phase of the project.

